# A new species of *Lethrinops* (Cichliformes: Cichlidae) from a Lake Malawi satellite lake, believed to be extinct in the wild

**DOI:** 10.1101/2023.03.17.533142

**Authors:** George F. Turner, Denise A. Crampton, Martin J. Genner

**Affiliations:** School of Natural Sciences, Bangor University, Bangor, Gwynedd LL57 2UW, United Kingdom & Vertebrates Division, Natural History Museum, Cromwell Road, London SW7, UK; School of Natural Sciences, Bangor University, Bangor, Gwynedd LL57 2UW, United Kingdom; School of Biological Sciences, University of Bristol, Life Sciences Building, 24 Tyndall Avenue, Bristol, BS8 1TQ, United Kingdom

**Keywords:** African cichlid, haplochromine, Lake Chilingali, morphology

## Abstract

A new species of cichlid fish, *Lethrinops chilingali* is described from specimens collected from Lake Chilingali, near Nkhotakota, Malawi. It is assigned to the genus *Lethrinops* based on the form of the lower jaw dental arcade and by the absence of traits diagnostic of the phenotypically similar *Ctenopharynx, Taeniolethrinops* and *Tramitichromis*. It also lacks the enlarged cephalic lateral line canal pores found in species of *Alticorpus* and *Aulonocara*. The presence of a broken horizontal stripe on the flanks of females and immature/non-territorial males of *Lethrinops chilingali* distinguishes them from all congeners, including *Lethrinops lethrinus*, in which the stripe is typically continuous. *Lethrinops chilingali* also has a relatively shorter snout, shorter lachrymal bone and less ventrally positioned mouth than *Lethrinops lethrinus*. It appears likely that *Lethrinops chilingali* is now extinct in the wild, as this narrow endemic species has not been positively recorded in the natural environment since 2009. Breeding populations remain in captivity.

## Introduction

Satellite lakes are small lakes lying in the catchment of much larger lakes, formerly or sometimes intermittently connected (Kaufman & Ochumba 1993; Mwanja et al. 2001; Genner et al. 2007). Their presence has been proposed to enhance the generation of biodiversity by isolating populations and facilitating allopatric speciation. Their role in the generation of African cichlid fish diversity was highlighted by the discovery of unique haplochromine cichlid fishes in Lake Nabugabo in the Lake Victoria catchment (Greenwood, 1965). Subsequently, several satellite lakes around Lake Malawi have also been shown to be inhabited by unique haplochromine cichlid fish populations (Turner *et al*., 2019). These satellite water bodies include Lake Chilingali, a small lake lying on the Kaombe River which flows into the middle of the western shoreline of Lake Malawi near Nkhotakota, from which a phenotypically distinct haplochromine species informally referred to as *Lethrinops* sp. “chilingali” (Tyers et al. 2014; Turner et al. 2019) has been sampled.

The genus *Lethrinops* Regan 1922 is currently used for haplochromine cichlids endemic to the Lake Malawi catchment distinguished by the semicircular shape of the dental arcade of the outer series of lower jaw teeth, which curves round to end abruptly behind the inner row(s), if present (Trewavas 1931, Turner 1996, Ngatunga & Snoeks 2004). This character is also found in the genera *Taeniolethrinops* Eccles & Trewavas 1989 and *Tramitichromis* Eccles & Trewavas 1989 which were split off from *Lethrinops* by Eccles & Trewavas (1989). The character is also known in a single species of the genus *Ctenopharynx* Eccles & Trewavas 1989 [*Ctenopharynx pictus* (Trewavas 1935)]. All of these taxa have ventrally positioned mouths, and relatively flat lower jaws with thin mandibular bones and small teeth. This jaw structure is believed to be associated with their feeding behaviour, which, where known, largely consists of ‘sediment-sifting’ or ‘winnowing’ (Weller *et al*. 2022), whereby loose sand or mud is picked up in the mouth, tumbled briefly and then ejected through the mouth and / or operculum, presumably with prey retained and swallowed (Fryer 1959; Fryer & Iles 1972; Konings 2016). Species in the genus *Lethrinops* are largely distinguished from *Taeniolethrinops, Tramitichromis* and *Ctenopharynx* by their lack of traits that distinguish those genera (Eccles & Trewavas 1989, Turner 2022). Not surprisingly, *Lethrinops* is currently believed to be polyphyletic (Ngatunga & Snoeks 2004; Malinsky et al. 2018; Masonick et al. 2022). Currently, the genus is ‘operational’, in the sense that it is possible to determine whether newly discovered taxa fall within its definition.

The purpose of the current work is to describe the Lake Chilingali species previously referred to as *Lethrinops* sp. ‘chilingali’ (Tyers et al. 2014; Turner et al. 2019) as *Lethrinops chilingali*, and to compare it with its presumed sister species from the main body of Lake Malawi, the morphologically similar *Lethrinops lethrinus* (Günther, 1893). The distributions of both species are discussed, and the current conservation status of *L. chilingali* is reviewed.

## Materials and methods

Specimens of the new species were obtained from fishermen on the shores of Lake Chilingali from 22-24 June 2009, euthanised with MS-222 (if still alive) and fixed in 10% formalin before being transferred to 70% alcohol (Industrial Methylated Spirit, IMS) for long term preservation. Additional specimens obtained from a captive strain kept at Bangor University euthanised in 2020 were preserved directly in IMS. These were used to investigate allometric comparisons between the two species, as they had grown to larger sizes than field-collected material. These captive bred fishes were excluded from the type series, but were included in statistical tests.

Comparative material of *L. lethrinus* included the holotype, and material from collections that were made in 1991-1992. These specimens were fixed in formalin and preserved in alcohol, along with some specimens collected in 2017 that were preserved directly in alcohol. Information on other congeneric species was obtained from literature, notably Trewavas (1931), Eccles & Lewis (1978), Eccles & Trewavas (1989), Turner (1996) and Ngatunga & Snoeks (2004). Counts and linear measurements were carried out following the methods of Snoeks (2004), and analysed using SPSS v27 (IBM, NY).

Geometric morphometric analyses were carried out on preserved specimens, photographed against a standard grey background with a scale for calibration. An initial tps file was constructed using image file names with tpsUtil v1.82 (Rohlf, 2015). A total of 15 landmarks (Figure 1) were then placed using tpsDig2 v2.32 (Rohlf, 2015): 1 anterior tip of upper jaw; 2 posterior tip of upper jaw (maxilla); 3-6 anterior, posterior, lower and upper point of eye; 7-8 beginning and end of dorsal fin; 9-10 beginning and end of anal fin; 11 anterior origin of pelvic fin; 12-13 lower and upper insertion of pectoral fin, 14 posterior margin of upper insertion of the operculum, 15 base of isthmus. The posterior of the caudal peduncle was not landmarked due to the upward flexion of the peduncle in several *L. lethrinus* specimens. Landmark data from the tps file were imported to MorphoJ v1.07 (Klingenberg 2011), where a Procrustes analysis was used to transpose, rotate and scale them into comparable Procrustes coordinates. These were analysed using SPSS v27 (IBM, NY).

**FIGURE 1.**
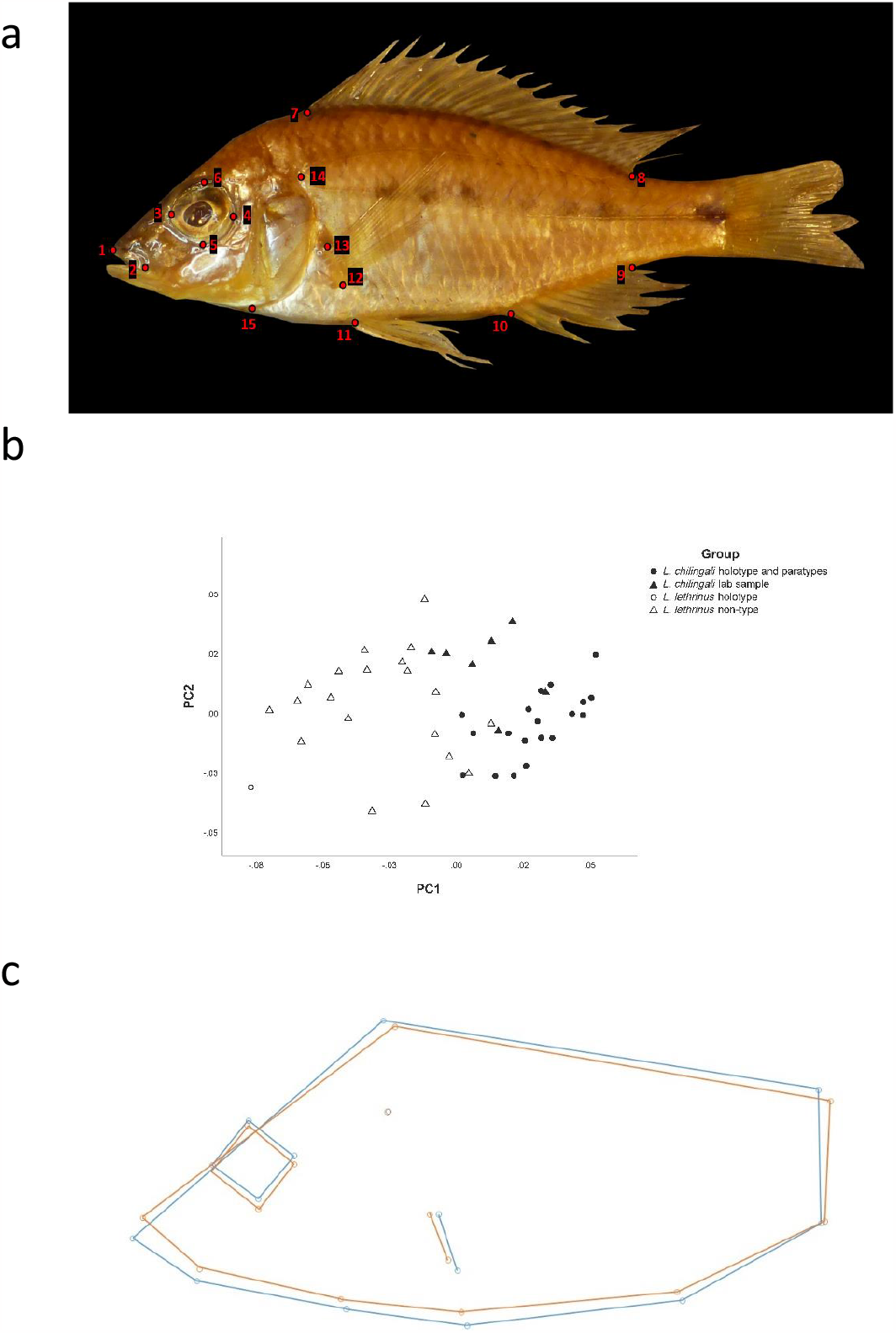
Geometric morphometric analyses of *Lethrinus lethrinus* and *Lethrinus chilingali* **a**. Landmarks used to quantify shape variation of preserved specimen (see Materials and methods for details). **b**. Principal Component Analysis indicates strong separation of *L. lethrinus* and *L. chilingali* on PC1, with clear differentiation of the respective holotypes. **c**. Wireframe plots of mean body shapes of *L. lethrinus* (blue) and *L. chilingali* (orange), showing the more ventrally placed mouth, longer snout, and higher back in specimens of *L. lethrinus* relative to *L. chilingali*

**FIGURE 2.**
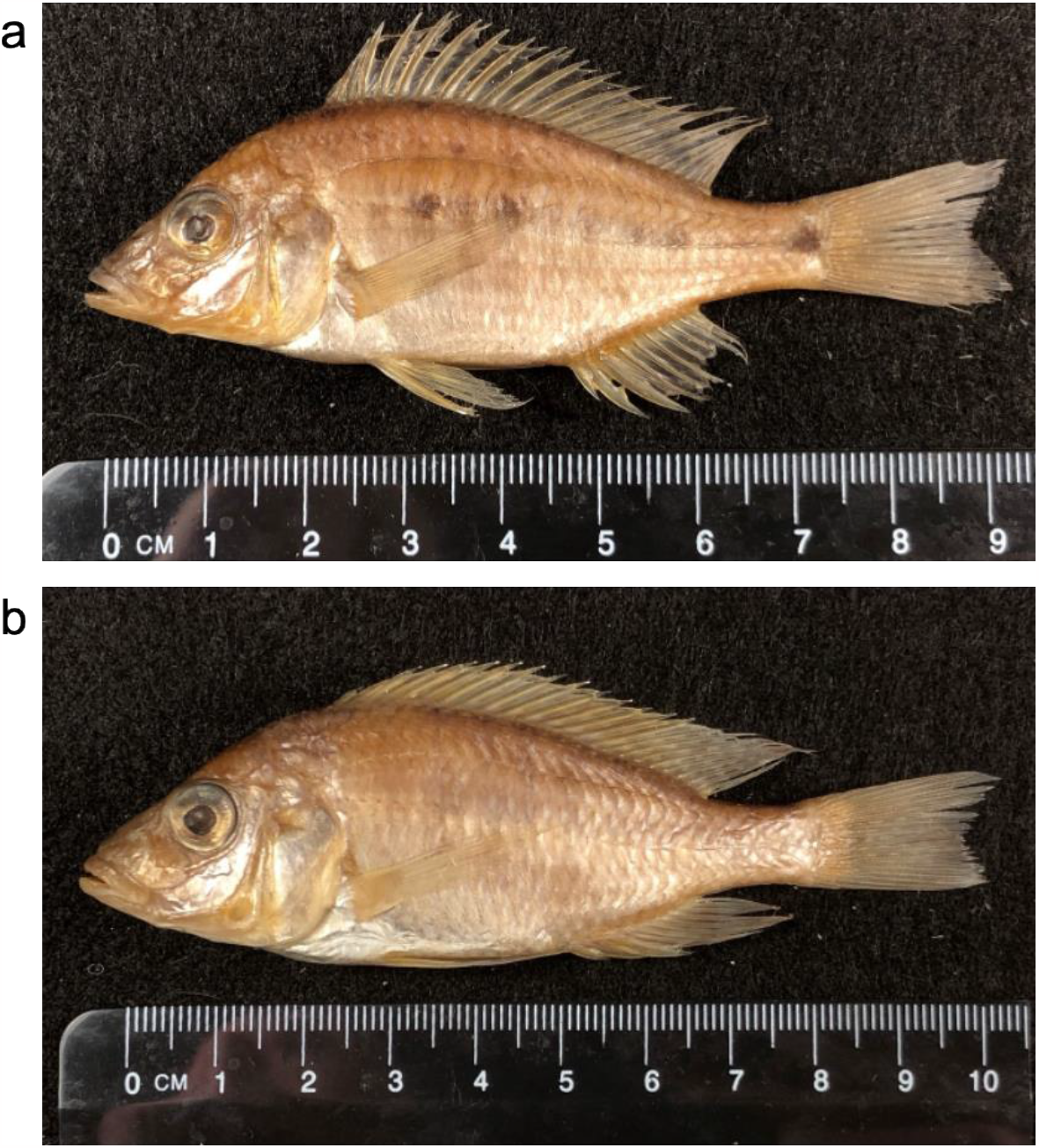
*Lethrinops chilingali*. **a**. Holotype, BMNH 2023.1.11.1; female 70.9mm SL. **b**. Paratype, BMNH 2023.1.11.2-21; mature male, 81.2mm SL.

Observations of live fish were collected from stocks descended from wild-caught fish obtained from Lake Chilingali between 2004 and 2009. Information on diets was taken from previous publications (Tyers et al. 2014; Turner et al. 2019), and an additional three wild-caught specimens of the new species were dissected to inspect stomach contents. Data and images used in analyses are available at: https://doi.org/10.5281/zenodo.8007304.

## Results

### Quantitative comparisons

Geometric morphometric data were ordinated using a Principal Component Analysis, with the primary axis (PC1) and secondary axis (PC2) capturing 34.2 and 19.6% of the variation, respectively. Overall, there was highly significant differentiation between *L. chilingali* and *L. lethrinus* on PC1 (General Linear Model; *F*_1,47_ = 39.25, *P* < 0.001), but not PC2 (*F*_1,47_ = 0.52, P = 0.60). The respective type specimens were among the most clearly differentiated individuals (Figure 1). The wireframe plots showed that the *L. lethrinus* specimens have a relatively more ventrally positioned mouth than *L. chilingali*, leading to a longer snout, and a deeper body at the anterior insertion of the dorsal fin.

Comparisons of linear morphometric measurements revealed significant differences in slopes of head length, anal fin base length and caudal peduncle length when regressed on standard length (Table 1). Assuming equal slopes, and using standard length as a covariable, *L. lethrinus* had significantly relatively greater body depth, interorbital width, snout length, lower jaw length, lachrymal bone depth, pre-pelvic length and caudal peduncle depth than *L. chilingali* (Table 1). The clearest differences were in snout length and lachrymal bone depth, followed by interorbital width.

**TABLE 1.** Comparison of linear morphometric measurements between *Lethrinops chilingali* (including captive-bred specimens) and *Lethrinops lethrinus* using General Linear Models and log_10_ transformed data. Bold indicates statistically significant differences between the species. * p < 0.05, ** p < 0.01, *** p < 0.001.

| Measurement | Slope |  | Elevation |  |
| --- | --- | --- | --- | --- |
| | $F_{1,51}$ | $P$ | $F_{1,52}$ | $P$ |
| Maximum body depth | 0.07 | 0.794 | **4.08** | **0.049\*** |
| Head length | **4.55** | **0.038\*** | 0.62 | 0.435 |
| Head width | 2.09 | 0.155 | 1.41 | 0.240 |
| Interorbital width | 1.73 | 0.198 | **10.80** | **0.002\*\*** |
| Snout length | 0.72 | 0.401 | **15.11** | **< 0.001\*\*\*** |
| Lower jaw length | 0.93 | 0.340 | 4.21 | 0.045 |
| Premaxillary pedicel length | 1.09 | 0.301 | 1.56 | 0.218 |
| Eye diameter | 1.02 | 0.317 | 1.79 | 0.187 |
| Lachrymal depth | 0.27 | 0.614 | **17.69** | **< 0.001\*\*\*** |
| Dorsal fin base length | 0.00 | 0.995 | 2.09 | 0.155 |
| Anal fin base length | **5.87** | **0.019\*** | 0.87 | 0.354 |
| Predorsal length | 3.42 | 0.070 | 0.17 | 0.686 |
| Preanal length | 1.20 | 0.279 | 0.06 | 0.815 |
| Prepectoral length | 0.06 | 0.808 | **5.41** | **0.024\*** |
| Prepelvic length | 1.08 | 0.303 | 2.51 | 0.119 |
| Caudal peduncle length | **4.96** | **0.030\*** | 0.92 | 0.341 |
| Caudal peduncle depth | 2.48 | 0.122 | **4.40** | **0.041\*** |

Comparisons of meristic counts showed that *L. lethrinus* had significantly more cheek scale rows than *L. chilingali* (K-S test, *Z* = 2.001, *P* = 0.001). Meanwhile *L. chilingali* had significantly more dorsal rays (K-S test, *Z* = 1.805, *P* = 0.003), upper gillrakers (K-S test, *Z* = 1.682, *P* = 0.007) and lower gillrakers (K-S test, *Z* = 2.903, *P* < 0.001) than *L. lethrinus* (Tables 2 & 3).There were no differences between the species in dorsal spines (K-S test, *Z* = 1.05, *P* = 0.221), anal spines (always 3), anal rays (K-S test, *Z* = 0.265, *P* ∼ 1.00) or lateral line scales (K-S test, *Z* = 0.98, *P* = 0.292) (Tables 2 & 3).

### *Lethrinops chilingali* new species

#### Holotype

BMNH 2023.1.11.1, female, 70.9 mm SL, collected from seine catches, Lake Chilingali (12.94°S, 34.21°E), 22-24 June 2009.

#### Paratypes

BMNH 2023.1.11.2-21, twenty specimens 59.3-81.2 mm SL, collected with holotype.

#### Other material (excluded from the type series)

BMNH 2023.1.11.22-28; seven specimens 56.8-98.7mm SL, laboratory bred from specimens collected at Lake Chilingali

#### Etymology

‘chilingali’ from Lake Chilingali, the type locality, used as a noun in apposition.

#### Diagnosis

The outer tooth row of the lower jaw curves smoothly to end just behind the inner tooth rows (*Lethrinops*-type dentition), distinguishing the species from other Lake Malawi haplochromines apart from species of the genera *Ctenopharynx, Lethrinops, Taeniolethrinops* or *Tramitichromis. Lethrinops chilingali* can be distinguished from other species in the genera *Ctenopharynx, Lethrinops, Taeniolethrinops* and *Tramitichromis* by the presence of a conspicuous horizontal series of dark grey to black spots along the middle of the flanks behind the head, linked to form one or two longer dashes, in total comprising 3-6 separate elements. *Lethrinops lethrinus* has a similar horizontal dark midlateral band, but it is typically more continuous, particularly posterior to the first anal spine, rather than broken into shorter spots and dashes. The horizontal melanic elements are generally not exhibited in dominant reproductively active males, however. *L. chilingali* also typically has a less ventrally placed mouth and shorter snout than *L. lethrinus* (snout as % of head length: 31.1-41.8 in *L. chilingali*, 37.6-50.0 in *L. lethrinus*).

#### Description

Body measurements and counts are presented in Table 2. *L. chilingali* is a small (<85mm SL in wild-caught specimens) moderately laterally compressed (maximum body depth 2.0-2.4 times maximum head width) cichlid fish with a moderately long snout (31.1-41.8 % head length). Females and immature males have distinctive melanic markings in the form of a horizontal row of dark spots and dashes (fig. 3b, d) and also have a thin red dorsal fin margin, while mature males are brilliant metallic green with a red dorsal fin margin above broader black and white bands (fig. 3f).

**TABLE 2.**
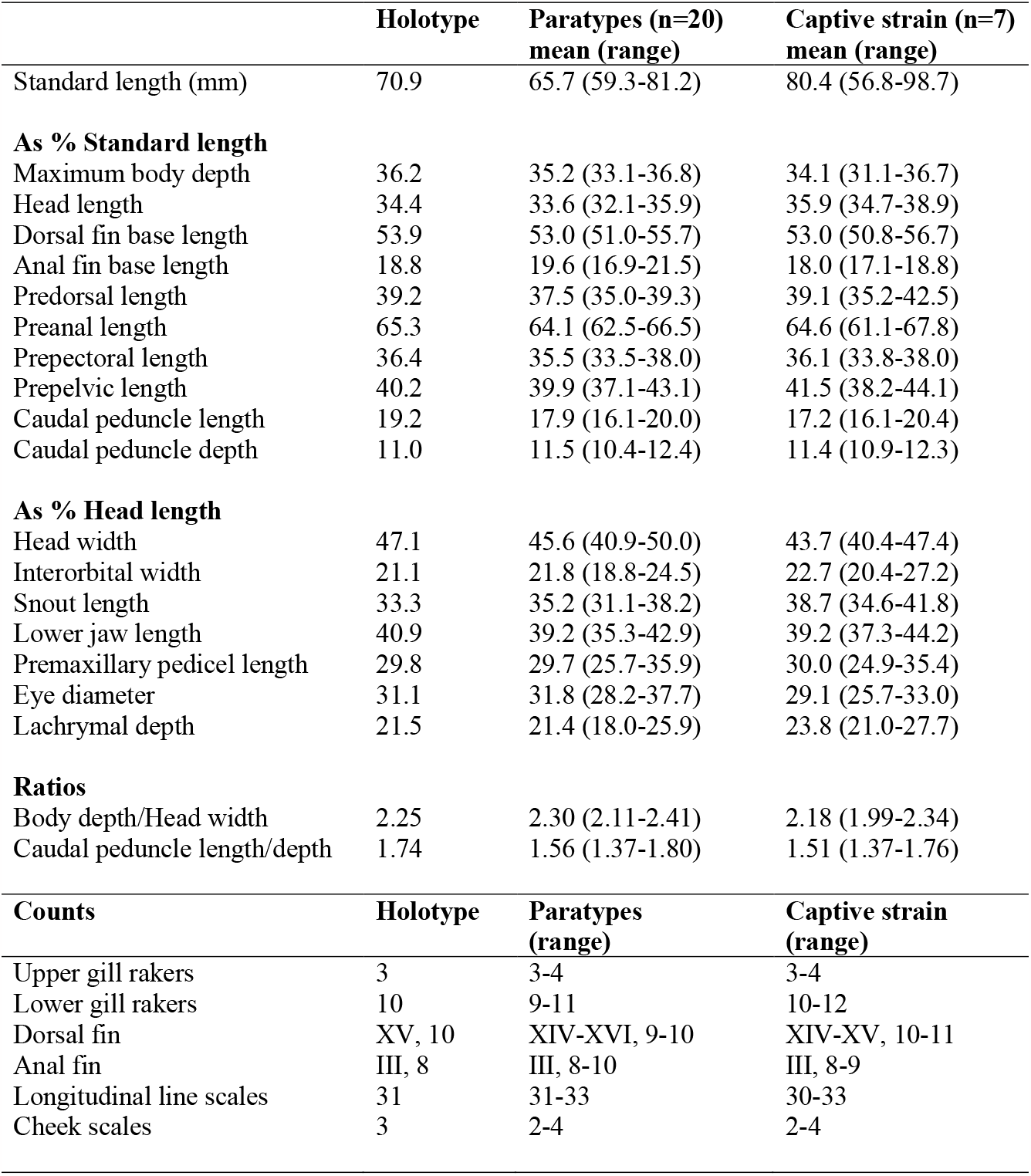
Morphometric and meristic characters of *Lethrinops chilingali*.

**FIGURE 3.**
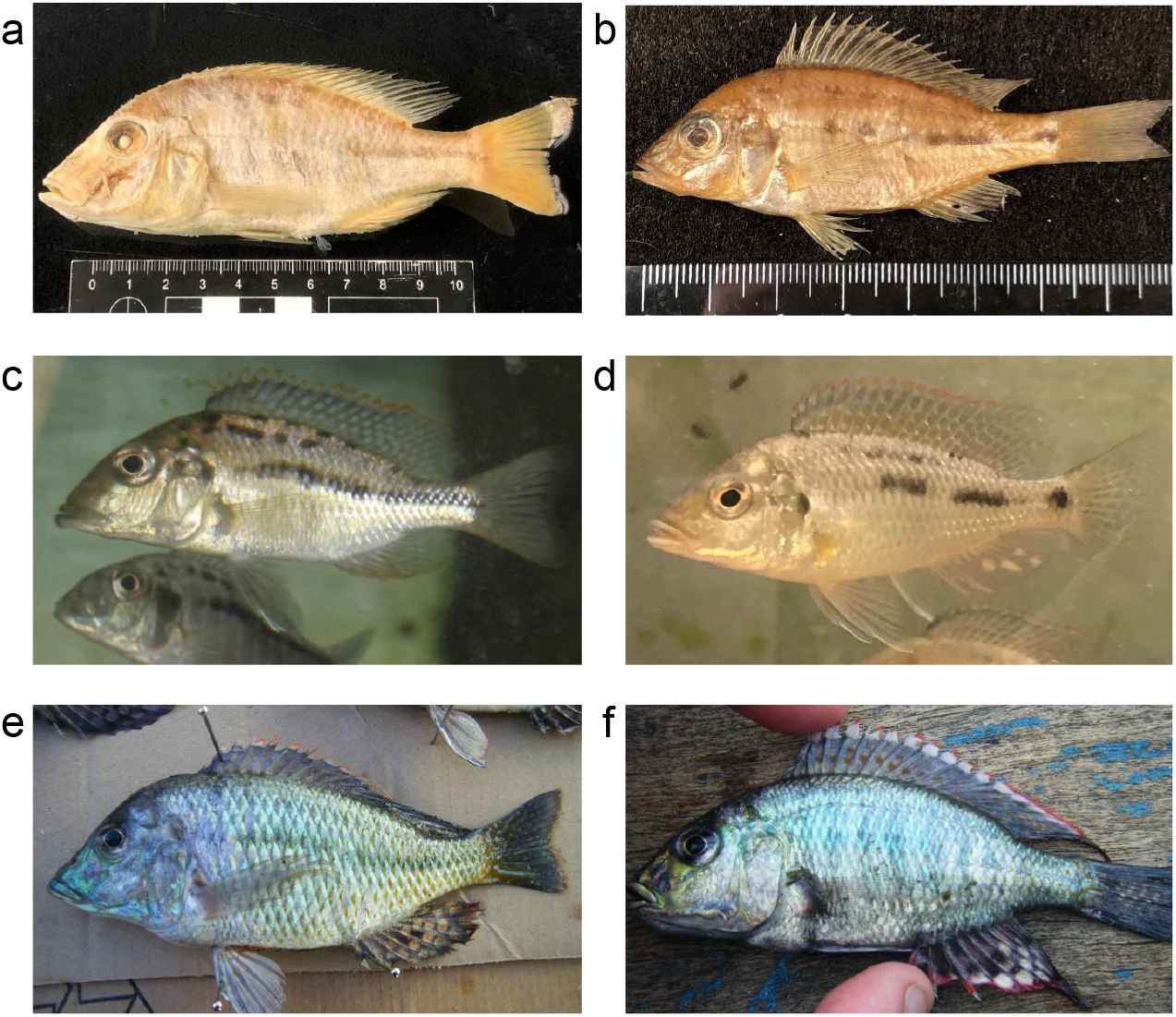
Comparisons of *Lethrinops lethrinus* and *Lethinops chilingali*. **a**. holotype of *L. lethrinus*, BMNH 1893.15.15., 118.5mm SL. **b**. paratype of *L. chilingali*, BMNH 2023.1.11.2-21, female, 60.7mm SL; **c**. *L. lethrinus* apparent female alive in aquarium. **d**. *L. chilingali* apparent immature male alive in aquarium. **e**. mature male *L. lethrinus*. **f**. mature male *L. chilingali*. The shorter snout *L. chilingali* is evident, and the more broken midlateral stripe can be seen in the live specimens.

All specimens are relatively deep-bodied and laterally compressed, with the deepest part of the body generally well behind the first dorsal fin spine. The anterior upper lateral profile is almost straight from the tip of the snout to the plane of the posterior margin of the eye, occasionally with a slight concavity above the eye, gentle sloping at an angle of about 40-degrees to the horizontal plane. There is no inflection to the angle of the profile above the eye (in contrast to *Tramitichromis* and *Tropheops* Trewavas 1984) and the premaxillary pedicel makes little or no interruption to the profile. The tip of the snout lies at about the same level in a horizontal plane as the upper margin of the insertion of the pectoral fin and at or below the level of the lowermost margin of the eye. The lower anterior lateral profile is also almost straight as far as the insertion of the pelvic fins, gently angled to the horizontal plane (about 12-15-degrees) and with little inflection at the posterior angle of the lower jaw even when the mouth is fully closed. The lower profile is more or less horizontal between the pelvic and anal fins. The mouth is relatively small and moderately upwardly-angled (gape about 40-degrees to horizontal). The caudal peduncle is relatively slender (1.4-1.8 times longer than deep). The pectoral fins are relatively long, extending past the first anal spine, but the pelvic fins are generally short of this, except in the largest mature males. The dorsal and anal fins, when folded, end well short of the caudal fin insertion, except in large mature males. The caudal fin is crescentic. The eye is large and circular and almost touches the upper lateral profile in perpendicular lateral view.

The flank scales are weakly ctenoid, with the ctenii becoming reduced dorsally, particularly anteriorly above the upper lateral line, where they transition into a cycloid state. The scales on the chest are relatively large and there is a gradual transition in size from the larger flank scales, as is typical in non-mbuna Lake Malawi endemic haplochromines (Eccles & Trewavas 1989). A few small scales are scattered on the proximal part of the caudal fin.

The cephalic lateral line pores are inconspicuous and the flank lateral line shows the usual cichlid pattern of separate upper and lower portions. The lachrymal bone is about as wide as deep and the lateral line pores are heavily overgrown with skin.

The lower jaw is relatively small, with thin mandibular bones. The jaw teeth are small, short and erect. The outer series in both the upper and lower jaw are short, blunt, erect and largely unequally bicuspid. These is a single inner series of small, pointed tricuspid teeth.

The lower pharyngeal bone (fig. 4a) is small, lightly built, Y-shaped, and carries small, slender, widely-spaced simple teeth, as illustrated for *L. lethrinus* by Eccles & Lewis (1978, figure 5). The teeth gradually increase in size from lateral to medial positions, but there are no distinctly differentiated enlarged medial teeth. There are approximately nine teeth in the midline row and 17-18 on each side on the posterior row. The gill rakers are short and blunt, generally with the most anterior rakers in the lower and upper arches reduced to small stubs.

**FIGURE 4.**
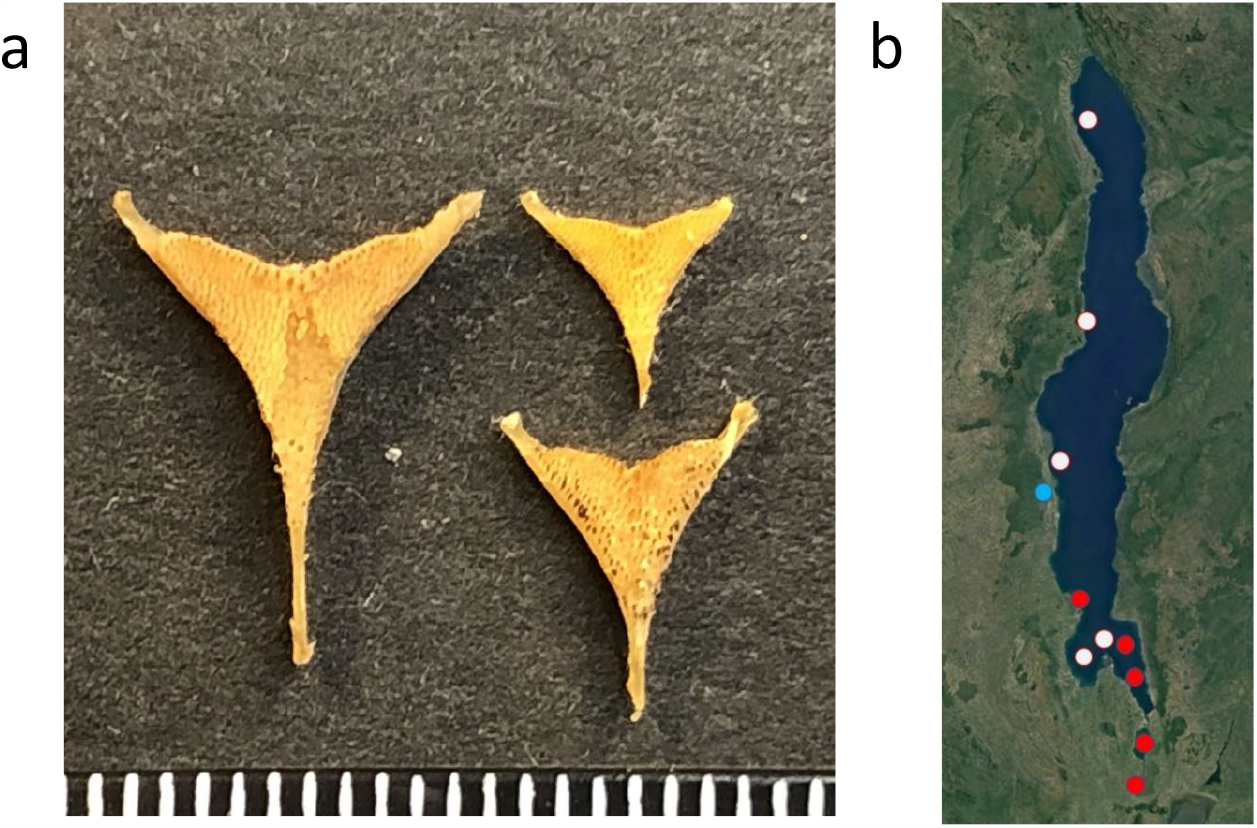
a. Lower pharyngeal bones of *Lethrinops lethrinus*, 128mm SL, BMNH 2023.1.11.49-50 (left); *Lethrinops lethrinus* 84mm SL (unregistered, bottom right); *Lethrinops chilingali* 69mmSL (unregistered, top right); b. Distribution of *Lethrinops lethrinus* specimens examined 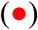, unconfirmed records: juveniles or not examined 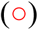 and *Lethrinops chilingali* 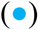.

**FIGURE 5.**
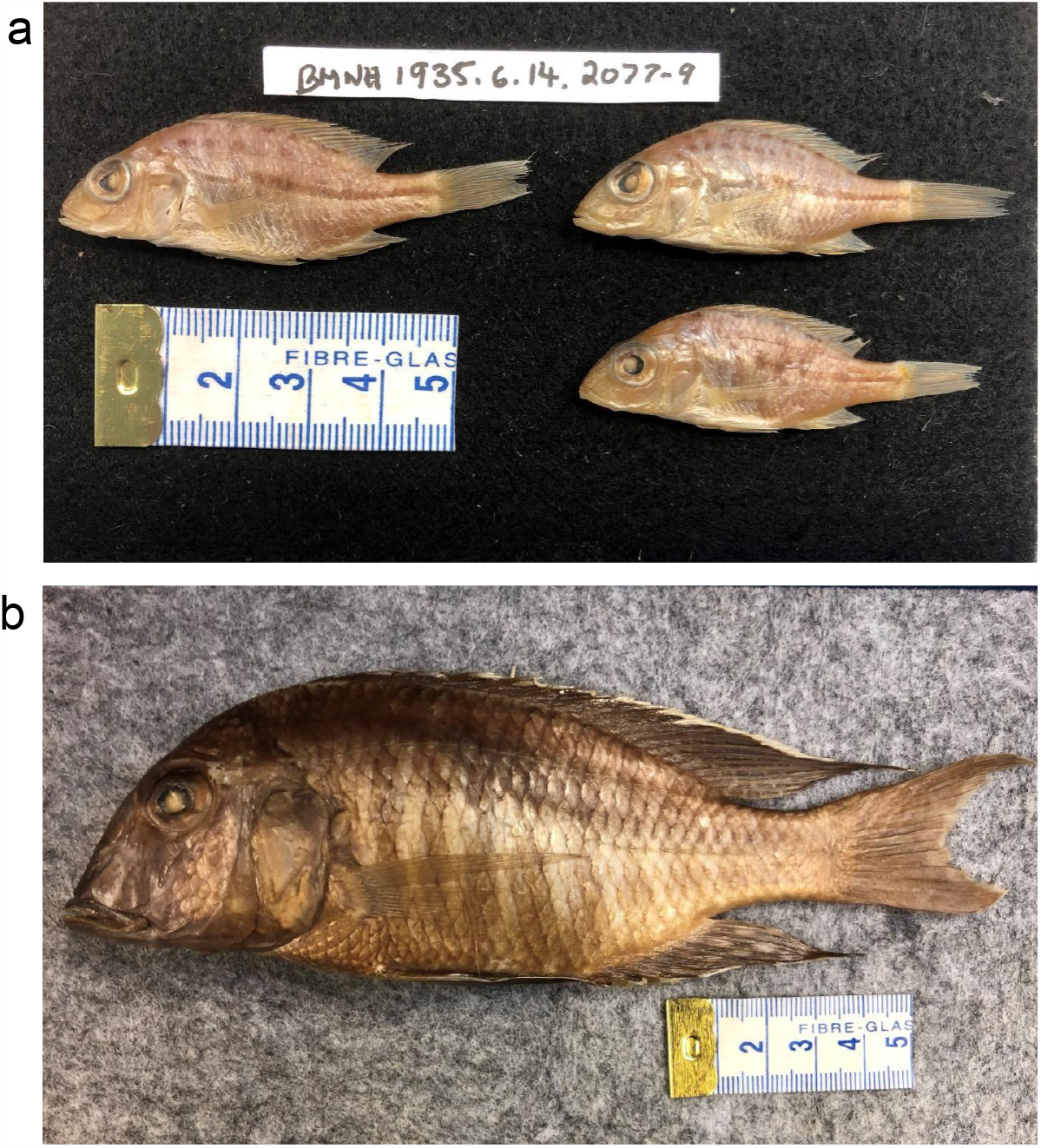
Comparative material. **a**. Three small specimens (BMNH 1935.6.14.2077-9) from Lupembe in northern Lake Malawi match *Lethrinops lethrinus*, in melanin pattern and low position of mouth on the head. **b**. A syntype of *Lethrinops leptodon* BMNH 1921.9.6.201-207, showing two oblique stripes thickened and fused together to form a midlateral blotch. This pattern is distinguishable from those of *L. chilingali* and *L. lethrinus*, but is similar to the Nkhata Bay population reported by Eccles & Lewis (1978) and assigned by them to *L. lethrinus*.

Female and immature fish (fig. 3d) are countershaded, pale sandy-brown dorsally, pale silvery on the flanks and underside. The flanks are marked by a midlateral horizontal row of dark spots and stripes extending from just behind the upper part of the operculum to the caudal peduncle. This varies between individuals, but generally comprises three to six separate melanic elements. A series up to six dark blotches is sometimes visible at the base of the dorsal fin, and element of a thin longitudinal dark stripe sometimes appears about half-way between the midlateral stripe and the base of the dorsal fin, usually starting a little behind the head and ending well before the caudal peduncle. The dorsal fin has a thin red outer margin and occasionally shows some faint dark spotting on both spinous and soft portions. Occasionally there is a pale submarginal band and anteriorly a thicker dark band. The caudal fin is usually translucent, sometimes with faint spotting. The anal fin sometimes shows a few faint yellowish spots.

Males in breeding dress (fig. 3f) are iridescent metallic green to pale blue. The horizontal melanic markings are occasionally exhibited when individuals are caught in fishing gear, or defeated in aggressive contests (seen in aquaria). Sometimes a series of faint dark vertical bars are visible. Patches of flank scales sometimes exhibit a metallic orange section anteriorly. The dorsal fin has a broad scarlet margin, underlain with a white submarginal band: these bands are narrower on the soft dorsal area. On the spinous dorsal, the red and white bands are underlain with a broad black band which extends to the base of the dorsal fin on the first inter-radial membrane, but as the membranes become longer posteriorly, the band overlies a series of orange spots extending onto the soft dorsal area, where they can be up to 10 spots between the longest rays. The membranes between the spots are pale grey to white. The caudal fin continues this pattern of orange spots with white/grey areas between. Sometimes the white inter-spot areas are missing, resulting in spots merging into stripes parallel to the fin rays. Occasionally, the white areas merge into stripes too. The upper and lower parts of the caudal fin can sometimes appear a bit darker, particularly on the basal section closer to the body, and particularly during dominant/courting behaviour. The pelvic fins are dark grey to black with a thin white anterior edge. The anal fin is greyish to black depending on mood, with a wide pink to red lower margin. A variable number (4-18) of large pale yellow ‘egg-spots’ are visible in one to two rows on the membranes behind the third anal spine. The colour of the iris varies from silvery to dark gold, with a darker spot above and below the lens continuing the line of a dark lachrymal stripe from the corner of the mouth. This stripe is very variable in intensity, showing up very prominently during territorial defence and courtship phases. The lower surface of the head and chest can turn dark grey during courtship and territorial behaviour but is otherwise pale greyish.

#### Distribution

Known only from Lake Chilingali in the Lake Malawi catchment (fig. 4b).

#### Behaviour and Ecology

The diet of *L. chilingali* specimens sampled in 2009 consisted largely of chaoborid (midge) larvae and pupae, along with cladocerans and other larger invertebrates, including odonatan nymphs and caridinid shrimps, but with little detritus, perhaps suggesting more midwater feeding than is usual in *Lethrinops* species. The behaviour of the species in the wild has not been observed, as the water of Lake Chilingali was highly turbid when visited between 2004 and 2009.

In captivity, *L. chilingali* females, non-territorial males and juveniles tend to aggregate in loose groups, feeding not only in the sediment, but on objects such as rocks or plants, or even at the surface. When attempts are made to catch the fish, they show a strong tendency to dive into the sand, turning sideways and completely burying themselves. This same behaviour has been reported to occur in the wild in *Fossorochromis rostratus* (Boulenger 1899), another cichlid from the Lake Malawi radiation (Fryer & Iles 1972, p. 207).

Dominant male *L. chilingali* are territorial and actively court females in typical haplochromine style: fins wide open, body horizontal or head-up, making rapid darts to the spawning site and back to the female, with spawning taking place amid bouts of circling and quivering, while alternating head-to-anal-fin ‘T-positions’ on the substrate. It is notable that dominant male coloration and aggression vary a lot, appearing to peak when females are approaching spawning, but are otherwise often quite subdued. During persistent bouts of courtship or aggression, the melanic elements of the male colour are emphasised, particularly the lachrymal/eye stripe, dark pelvic and anal fins, dark upper and lower margins of the caudal fin and even faint vertical barring on the flanks. Even in a large tank with a high density of fish, there is usually just a single dominant male: this is similar to *Astatotilapia* Pellegrin 1904, which tend to be solitary breeders. Communal lek breeders, such as *Oreochromis* Günther 1889 will usually divide up a tank into numerous smaller territories and engage in frequent boundary disputes. This suggests that *Lethrinops chilingali* are not communal lek breeders in the wild.

There is little indication of bower construction in *L. chilingali* when a sand or gravel substrate is provided: dominant males usually try to lead females to a slight depression near to an object such as a rock or piece of wood: in a bare tank, the focus would probably be the tank bottom near one of the corners or a wall near a heater or filter inlet. This is in marked contrast to reports of *L. lethrinus* where complex bowers have been recorded in the field, out over open substrate (Konings 2016, p. 369). In *L. chilingali*, the construction of the depression seems almost haphazard: males have not been observed to show consistent bouts of digging, but spend most of their time chasing, then returning to the territory focus next to the object, during which they make occasional ‘feeding movement’ of picking up a mouthful of substrate, moving forwards and ejecting it through the mouth and/or opercular openings at a slight distance away. This occurs all over the vicinity of the side of the object they are defending, but there seems to be a slight bias towards a certain point up against the object, which thereby becomes a shallow depression.

Female *L. chilingali* are maternal mouthbrooders, brooding young until they are capable of independent feeding. As fry complete the absorption of the yolk, they show through the female’s buccal membrane as a dark area, but females do not develop the ‘warpaint’ typical of fry guarders, such as known *Astatotilapia* or *Oreochromis:* dark eyes, lachrymal stripes and forehead stripes. There is no indication that females guard free-swimming fry or permit them to return to their mouths. This non-guarding behaviour is similar to other known shallow-water *Lethrinops* species.

### *Lethrinops lethrinus* (Günther, 1893)

#### Holotype

*Lethrinops lethrinus* (Günther, 1893): BMNH 1893.11.15.15, 116.1 mm SL, coll. A. Whyte, Upper Shire River at Fort Johnston (Mangochi), March 1892,

#### Other material examined

BMNH 2023.1.11.29, 1 specimen 130.1mm SL, collected by G.F. Turner from experimental trawl at depth of 5-18m, between Namiasi and Palm Beach (approximately 14.38°S, 35.22°E), SE Arm of Lake Malawi, 30 July 1991.

BMNH 2023.1.11.30, 1 specimen, 120.6 mm SL, collected by G.F. Turner, trawled at 5-18m depth between Namiasi and Malindi (approximately 14.34°S, 35.22°E), SE Arm of Lake Malawi, 30^th^ July 1991.

BMNH 2023.1.11.31, 1 specimen, 101.4mm SL, collected by G.F. Turner from kambuzi seine fisherman, West shore of Lake Malombe, probably at Chimwala (14.64°S, 35.18°E), 26 June 1992,

BMNH 2023.1.11.32-36, 5 specimens, 63.2-66.6 mm SL, collected by G.F. Turner from Lake Malombe, probably at Chimwala (14.64°S, 35.18°E), 23 July 1992.

BMNH 2023.1.11.37, 1 specimen 90.2 mm SL, collected by G. F. Turner, Middle Shire River, probably at Liwonde Barrage (15.06°S, 35.22°E), 20^th^ May 1992.

BMNH 2023.1.11.38-43, 6 specimens 129.2-152.6 mm SL, collected by G. F. Turner unspecified sites in southern Lake Malawi, 1990-1992.

BMNH 2023.1.11.44-46, 3 specimens 97.9-116.1 mm SL, collected by G. F. Turner, trawled at 18-21m at Ulande 1a station (14.23°S, 35.21°E), SE Arm Lake Malawi, 1991.

BMNH 2023.1.11.47-48, 2 specimens 106.4-130.2 mm SL, collected by David Bavin, from seine fishermen, Lake Malombe (14.64°S, 35.18°E), 6^th^ July 2009.

BMNH 2023.1.11.49-50, 2 specimens 121.7-128.0 mm SL, collected by G. F. Turner, trawled at 26m depth at Michesi station (14.32°S, 35.19°E), SE Arm of Lake Malawi, 1992.

BMNH 2023.1.11.51-53, 3 specimens 109.4-123.2 mm SL, collected by G. F. Turner, from seine net fishermen, Palm Beach (14.41°S, 35.23°E), SE Arm of Lake Malawi, 23 Jan 2017.

BMNH 2023.1.11.54, 1 specimen 120.7 mm SL, collected by G. F. Turner, from seine net fishermen, Palm Beach (14.41°S, 35.23°E), SE Arm of Lake Malawi, 22 Jan 2017.

#### Remarks

*L. lethrinus* was selected as the type of the genus *Lethrinops* by Regan (1922). It was originally described as *Chromis lethrinus* from a single specimen, but was redescribed from additional material by Regan (1922), Trewavas (1931), Eccles & Lewis (1978) and Eccles & Trewavas (1989). It was also included in a key to the shallow-water *Lethrinops* species by Ngatunga & Snoeks (2004). The original illustration in Günther (1893) shows a specimen with a continuous horizontal midlateral stripe beginning at the eye and extending to the base of the caudal fin. This is reprinted in Eccles & Trewavas (1989), where the imaged specimen is erroneously referred to as the lectotype (it is the holotype). The redescription by Eccles & Lewis (1978) includes a drawing of a non-type specimen in which the horizontal midlateral stripe is composed of a series of about 15 spots running from just behind the origin of the pelvic fin to the base of the caudal fin. Anteriorly, the first five spots are separate, but the gaps between them are much narrower than the length of the spots. Posteriorly, all of the spots overlap, to form a continuous, albeit irregular, blotchy line. Eccles & Lewis (1978) stated they examined (but did not measure) the type and there seems little doubt that the non-type material they studied (uncatalogued, Monkey Bay Fisheries Research Unit, Malawi, status unknown) corresponds to this species.

*Lethrinops lethrinus* is readily diagnosed based on its typical *Lethrinops*-type dentition, horizontal melanic flank markings and long snout. Mature males show a metallic blue-green breeding dress, with a prominent red and white dorsal fin margin and numerous large eggspots on the anal fin (Figure 3, see also Konings 2016). *L. lethrinus* appears to be confined to shallow waters with muddy bottoms, often river mouths with extensive beds of reeds and other macrophytes, feeding on invertebrates and other edible material obtained from the sediment (Turner 1996). Konings (2016) reports a lakewide distribution and it has been recorded from Lake Malombe and the Upper and Middle Shire Rivers (Turner 1996), but records from Domira Bay northwards are based on juveniles that are hard to distinguish from *L. chilingali* or lack available voucher material (fig. 4b). Counts and measurements of the material we examined are presented on Table 3.

**TABLE 3.** Morphometric and meristic characters of *Lethrinops lethrinus*.

|  | **Holotype** | **Non-types (n=26)**<br>**mean (range)** |
| --- | --- | --- |
| Standard Length (mm) | 118.5 | 110.7 (62.9-152.6) |
| **As % Standard length** |  |  |
| Maximum body depth | 36.4 | 37.2 (33.0-41.0) |
| Head length | 34.5 | 35.1 (33.1-39.1) |
| Dorsal fin base length | 53.9 | 53.4 (49.9-56.2) |
| Anal fin base length | 17.7 | 18.7 (16.5-21.0) |
| Predorsal length | 37.7 | 39.5 (37.1-42.3) |
| Preanal length | 66.2 | 64.9 (61.5-68.6) |
| Prepectoral length | 35.1 | 37.1 (33.9-40.1) |
| Prepelvic length | 42.5 | 42.2 (35.7-46.4) |
| Caudal peduncle length | 17.9 | 17.5 (14.7-20.2) |
| Caudal peduncle depth | 12.5 | 12.1 (10.8-13.4) |
| **As % Head length** |  |  |
| Head width | 46.2 | 44.8 (41.0-50.1) |
| Interorbital width | 24.2 | 22.6 (18.0-26.9) |
| Snout length | 42.5 | 44.4 (37.6-50.0) |
| Lower jaw length | 40.1 | 41.0 (37.0-43.5) |
| Premaxillary pedicel length | 31.1 | 31.0 (25.4-34.3) |
| Eye diameter | 29.5 | 28.4 (25.2-34.7) |
| Lachrymal depth | 29.5 | 30.7 (21.6-34.8) |
| **Ratios** |  |  |
| Body depth/Head width | 2.29 | 2.36 (2.14-2.67) |
| Caudal peduncle length/depth | 1.43 | 1.45 (1.17-1.66) |
| **Counts** | **Holotype** | **Non-types (range)** |
| Upper gill rakers | 3 | 2-4 |
| Lower gill rakers | 9 | 8-10 |
| Dorsal fin | XV, 11 | XIV-XVI, 8-12 |
| Anal fin | III, 9 | III, 8-9 |
| Longitudinal line scales | 31 | 30-36 |
| Cheek scales | 3 | 3-4 |

## 4. DISCUSSION

### Relationship of *L. chilingali* to other taxa in the Lake Malawi radiation

The present study has assumed that *L. lethrinus* is both the most likely sister taxon for *L. chilingali* and the species most likely to interbreed with it, should habitat barriers be broken down. The former proposition is based on their overall similar appearance, including very similar male breeding dress, and similar – although distinct-melanin patterns in the females and juveniles. They are the only two known *Lethrinops* species to share a largely horizontally-banded melanin pattern. Other Lake Malawi cichlids also share some of these features, notably species of *Protomelas* Eccles & Trewavas 1989 found in similar shallow weedy/muddy habitats, including *Protomelas kirkii* (Günther 1894), *Protomelas similis* (Regan 1922) and *Protomelas labridens* (Trewavas 1935) (Eccles & Trewavas 1989, Konings 2016, Turner 1996). These three species also have females/immatures with a sandy/countershaded appearance, with a strong horizontal dark band running along the flank. Males are also metallic blue-green, with a red and white dorsal fin margin. These species have shorter snouts and more upwardly-angled mouths than *L. lethrinus*, but so does *L. chilingali*, which is arguably morphologically intermediate between them. The genera *Protomelas* and *Lethrinops* can be distinguished by the shape of the lower jaw dental arcade, and it is presently assumed that this is a phylogenetically informative trait (Eccles & Trewavas 1989), although this requires confirmation from a phylogenetic analysis, ideally based on genome-scale sequence data. A published phylogenomic analysis places *L. lethrinus* in the middle of a clade of shallow water *Lethrinops, Taeniolethrinops* and *Tramitichromis* (Masonick et al. 2022), thus grouping these genera showing *Lethrinops*-type dentition (Eccles & Trewavas 1989). However, *P. kirkii, P. similis* and *P. labridens* were not included in that analysis (Masonick et al. 2022). Notably, however, an additional group of deep-water *Lethrinops* appears in a separate part of the phylogenetic tree, suggesting that the *Lethrinops*-type dentition is prone to parallelism. Thus, we conclude that available evidence does not conflict with *L. chilingali* being a sister species to *L. lethrinus*, but this requires detailed phylogenetic investigation for confirmation. If *L. lethrinus* shows relatively high levels of population structure, it could be paraphyletic (ancestral) with respect to *L. chilingali*.

### Distributions of *L. chilingali* and *L. lethrinus*

*Lethrinops chilingali* has only been positively recorded from Lake Chilingali, but here we consider whether it may have a broader distribution in Lake Malawi, possibly extending to the central to northern part of the lake as an allopatric sister species to *L. lethrinus*. Although a lake-wide distribution has been claimed for *L. lethrinus* (Konings 2016), the great majority of records backed by preserved specimens or photographs come from the southern arms, Lake Malombe and the Shire River (Eccles & Lewis 1978, Turner 1996, Konings 2016). On the Global Biodiversity Information Facility website (GBIF 2023), there is a record of *Lethrinops lethrinus* from co-ordinates indicating a collection site off the Tanzanian shore near Ngkuyo Island, Mbamba Bay (11.334°S, 34.769°E), based on specimens at the South African Institute for Aquatic Biodiversity (SAIAB). An offshore location near a rocky headland seems an unlikely collecting site for *Lethrinops lethrinus*, which favours shallow sheltered vegetated habitats and the locality label is given as ‘Lifuwu’, which probably corresponds to the vicinity of Lifuwu village (13.69°S, 34.60°E) just north of Salima, suggesting that the co-ordinates have been recorded in error. The single small specimen shows no melanic markings (faded post-preservation?), but the head shape is consistent with *Lethrinops lethrinus* rather than *L. chilingali*. Another GBIF record from co-ordinates 13.35°S, 33.4°E would suggest specimens were collected from the Rusa River, a tributary of the Bua River, which joins Lake Malawi near Lake Chilingali. The site is far upstream, around 97km West of the Lake Malawi shore at Benga, and initially we thought this might suggest specimens of *L. chilingali* could be widespread in the river catchment. However, the collection label indicates the specimens were collected from Lake Malawi at Foo, which is a trawling station in the SE Arm of Lake Malawi (also sometimes written as Fowo), which is at approximately 14.14°S, 35.18°E, again suggesting an error in the co-ordinates. Photographs of the specimens show typical *Lethrinops lethrinus*, with long snouts and strong horizontal melanic markings. The catalogue of the Natural History Museum in London contains a single accession of three specimens labelled as *L. lethrinus* from Lupembe Sand Bar, collected by Cuthbert Christy in 1925 (BMNH 1935.6.14.2077-9; Figure 4). The electronic catalogue suggests that this location is in Tanzania, perhaps following Eccles & Trewavas (1989) who suggested it might represent a site at the mouth of the ‘Kivira River’. However, the town at the mouth of the Kiwira River (as presently named) is currently known as Itungi Port. It is more likely that the 1925 collection site is Lupembe on the Malawian lakeshore, just south of Karonga (10.055°S, 33.99°E). Notably, recent satellite images show a conspicuous sandbar (Google Earth). Examination of the unpublished diary of Cuthbert Christy held at the Natural History Museum shows a single accession from Lupembe following an extensive collection of several hundred accessions from Vua / Deep Bay (Chilumba area) and immediately before another extensive collection from Mwaya in Tanzania, on the far north coast of the lake (itemising various river mouths visited). No other accessions were made from Lupembe. This suggests that the location was visited en-route from Chilumba to Tanzania, which would fit well with the location near Karonga. Unfortunately, the specimens (fig. 5a) are very small (44.8-50.9 mm SL) which makes morphological comparisons difficult with the larger specimens examined for this study, due to allometric effects. For example, they have notably relatively large eyes, making snout measurements relatively small. However, the low position of the mouth on the head and the largely continuous midlateral stripe, fit far better with *L. lethrinus* than with *L. chilingali*. Thus, available museum specimens strongly support the occurrence of typical *Lethrinops lethrinus* only in the southern arms of the lake, but tentatively indicate that they may also occur just north of Senga Bay and indeed almost at the northernmost extremity of the lake, but do not provide evidence for the occurrence of *L. chilingali* or any other similar form within Lake Malawi,

Other published records are not backed by specimens available to examine or photographic evidence. Eccles & Lewis (1978) reported that they had found *L. lethrinus* at Nkhata Bay, which is well to the north of Lake Chilingali. However, they reported an unusual melanin pattern: “the dark line along the middle of the flank curves upwards anteriorly to merge with the lower of the two rows of spots and the spots themselves may run together posteriorly to form a stripe”. The occurrence of specimens with dramatically different stripe patterns at Nkhata Bay might lend credence to the idea that *L. lethrinus* represents a complex of allopatric taxa, which might increase the probability that *L. chilingali* might persist in the main Lake Malawi. Eccles & Lewis provided no illustration of this ‘Nkhata Bay variant’. Their specimens were deposited in the collection of the Monkey Bay Fisheries Research Unit, Malawi and their present status is unknown. The pattern described is reminiscent of that seen on some of the type specimens of *L. leptodon* Regan 1922 (fig. 5b). In the same 1978 paper, Eccles & Lewis redescribed that species based on a single specimen collected from Chintheche in the NW of the lake, near Nkhata Bay, but their illustration of that specimen showed a clear midlateral blotch on the upper part of the flank. They reported examining, but not measuring, three of the type specimens of *L. leptodon*, which are held at the Natural History Museum in London (BMNH 1921.9.6.201-207, collected by Wood from somewhere in ‘Lake Nyasa’). Thus, it seems unclear whether the reported Nkhata Bay populations represent *L. lethrinus* or *L. leptodon*, or indeed something else. In summary, the status of the northern populations of *Lethrinops* of this group is unclear but is consistent with the hypothesis that *L. lethrinus* is found in suitable habitats throughout Lake Malawi, and that *L. chilingali* is a satellite lake endemic extinct in the wild.

### Conservation status of *Lethrinops chilingali*

Lake Chilingali is approximately 5km in length and a maximum of 1km in width, and is characterised by two deeper basins of approximately 5m depth separated by a shallower plateau (Turner et al. 2019). It has a single outflow, the Kaombe River, which meanders for approximately 12km before reaching the main body of Lake Malawi (Genner et al. 2007). The lake is a natural water body, and the two basins of the modern lake are represented on early European exploration maps, as two separate bodies of water, Lake Chikukutu to the south, and Lake Chilingali to the north (Turner et al. 2019). The lake level was raised when a dam was constructed across the single outflow for irrigation purposes, initially in the 1950s, before being modified in the early 1970s (Denys et al. 2013). The dam collapsed early in 2012 (Denys et al. 2013), and the single lake disappeared, reforming the two separate smaller shallow basins. In 2016 these basins were estimated to be only ∼1m deep and fringed with macrophytes. The lake was refilled to approximately its pre-collapse-level in June-July 2019 following the construction of a new dam.

During the period 2004 to 2011, before the collapse of the dam, *L. chilingali* was periodically and reliably sampled from the lake, alongside another apparently endemic haplochromine cichlid, the undescribed *Rhamphochromis* sp. “chilingali” (Genner et al. 2007; Turner et al. 2019). To our knowledge, the last sampling event where *L. chilingali* was recorded in the wild was on 25 June 2009 (by G. Turner), while representatives of *R*. sp. “chilingali” were last collected from an artisanal fishing catch on 12 January 2011 (by M. Genner). During sampling in February 2016, neither of the species was encountered in a survey of the main northern and southern basins of Lake Chilingali (Turner et al. 2019). A survey in April 2022 also failed to sample any either *L. chilingali* or *R*. sp “chilingali” but did find that Lake Malawi endemic *Otopharynx tetrastigma* (Günther 1894) was abundant (H. Svardal, pers comm). This species was absent between 2004 and 2016 and is likely to have been introduced during restocking after the lake was refilled in 2019 (H. Svardal, pers comm). Although further surveys of Lake Chilingali and the Kaombe river are warranted to determine if remnant populations of either *L. chilingali* or *R*. sp “chilingali” persist, on the basis of the current evidence, we consider it most likely that both species are no longer present in the natural environment. Breeding populations of *L. chilingali* or *R*. sp “chilingali” are, however, maintained in captivity, and may be candidates for reintroduction. On the basis of the evidence discussed above, we recommend that *L. chilingali* is attributed the status of Extinct in the Wild (EW) on the International Union for Conservation of Nature (IUCN) Red List of Threatened Species.

## Acknowledgements

We are grateful to the Malawi Government Department of Fisheries for collaboration and permits. Sampling and specimen collection on Lake Chilingali in 2004 was funded by the Natural Environment Research Council award NER/A/S/2003/00362, and in 2009 by a student expedition grant from Zoological Society of London to Gavan Cooke, Dave Bavin, Lucy Ferris, Cat Griggs and Bev Stubbs, to whom we are grateful for help in specimen collection. We thank Rupert Collins, Oliver Crimmen, James Maclaine and Simon Loader at the Natural History Museum in London for helping us with access to specimens, finding old field notes and cataloguing new material. We are grateful to Jay Stauffer and Roger Bills for photos of the SAIAB specimens and their labels, to Hannes Svardal for information about the 2022 expedition and to Alexandra Tyers and Dave Bavin for photographs of *Lethrinops lethrinus* in the aquarium and field respectively.

## Notes

### Competing Interest Statement

The authors have declared no competing interest.

### Summary of Updates

NB: new map and photos of Lower Pharyngeal bones, more details on the dam collapse, link to archived data, among other changes

